# Literacy acquisition and functional connectivity drive the emergence and continued development of the VWFA

**DOI:** 10.64898/2026.06.24.732877

**Authors:** Jin Li, Kelly J. Hiersche, Nii-Ayi Aryeetey, Anna Quatrale, Patricia Resnick, Zeynep Saygin

**Affiliations:** School of Psychology, Georgia Institute of Technology, Atlanta, GA, USA; Department of Psychology, The Ohio State University, Columbus, OH, USA; Center for Cognitive and Behavioral Brain Imaging, The Ohio State University, Columbus, OH, USA; Psychology, Georgia State University, Atlanta, GA, USA

**Keywords:** visual word form area, development, longitudinal, neuronal recycling hypothesis, connectivity hypothesis, fMRI

## Abstract

The visual word form area (VWFA) is a hallmark of literacy in the human brain. Alongside reading acquisition, the functional organization of the ventral temporal cortex (VTC) undergoes substantial changes. However, it remains unclear what factors uniquely drive the emergence and continued development of the VWFA, or more broadly shaping category selectivity across the VTC. Here, we combined cross-sectional and longitudinal data from children in early childhood (3-9 years) to investigate the development of visual language selectivity. We found that after controlling for age, literacy acquisition drove increases in word selectivity of the VWFA, but not the continued development of other early-developed category selectivity. Sensitivity to spoken higher-level language information within the VWFA was also associated with reading ability. By projecting the VWFA defined at the later time point onto each child’s earlier time point, we also found that the pre-VWFA showed no preferential tuning to any particular visual category. Finally, longitudinal changes within the VWFA, both increased responses to visually presented words and decreased responses to auditory control conditions, were associated with changes in functional connectivity of VWFA to high-level language regions, even after controlling for initial activity levels. Together, the current study shows a unique role for literacy acquisition and experience-dependent connectivity changes in the emergence and functional specialization of the VWFA, providing empirical evidence for the revised neuronal recycling hypothesis and connectivity hypothesis of functional brain organization.

## Introduction

Investigating the emergence of neural representations is crucial in developing a richer understanding of the factors that shape functional brain organization. The mosaic-like functional specializations to various high-level visual categories of the ventral temporal cortex (VTC) in adults (e.g., Kanwisher, 2010) are observed early in development; preferential responses to faces, scenes, and bodies are detectable in infants as young as three months (Deen et al., 2017; Kosakowski et al., 2022) but this functional specificity continues to refine over time (e.g., prolonged development of face and limb specialization) (Cohnen et al., 2026; de Haan et al., 2002; Golarai et al., 2007; Nordt et al., 2021; Taylor et al., 2004), suggesting an innate drive for initial representations, followed by continued neural tuning based on experience with these visual categories. Interestingly, unlike these evolutionarily important stimuli, the human VTC also houses a region dedicated to visual processing of written scripts, the visual word form area (VWFA). This word-selective region emerges only after a child learns to read (e.g., Baker et al., 2007; Cohen et al., 2002) but once it does emerge, it is located in roughly the same left-lateralized region across literate individuals. Therefore, examining the development of the VWFA allows researchers to test different theories on how the brain adapts to develop new selective regions. Critically, it remains unclear how neural tuning of the VTC to visual stimuli (including visual words) and to spoken language changes concurrently with, or as a consequence of, gaining literacy, and how increasing communication between VTC and high-level language cortex may contribute to these neural changes.

The neuronal recycling hypothesis is an appealing theory that proposes that specialization for written text is repurposed from functions that were originally devoted to other stimuli (Dehaene & Cohen, 2007), likely from those that require similar visual processing, such as foveal processing for faces (Behrmann & Plaut, 2015; Dehaene et al., 2010; Gomez et al., 2018). However, empirical evidence has failed to yield a clear picture: while competition between faces and words has been observed when comparing literate vs. illiterate adults (Dehaene et al., 2010), both cross-sectional and longitudinal developmental studies showed that face or object activity was not diminished, or invaded by newly developed word selectivity (e.g., van Paridon et al., 2021; Yeatman et al., 2024; see E. Kubota, Grotheer, et al., 2023 for a review). Revising the original theory, recent longitudinal studies have shown that word-selective responses are more likely to emerge from previously unspecified cortex that might show some weak sensitivity not to faces, but to other categories, like tools (Dehaene-Lambertz et al., 2018) and limbs (Nordt et al., 2021). However, while these longitudinal studies have provided valuable evidence for the developmental changes in other forms of category selectivity alongside the emergence of word selectivity, there is still a lack of evidence characterizing the response profile and neural changes within the pre-VWFA (the cortical tissue that will later develop as the VWFA) prior to reading, and directly linking these changes to children’s reading ability. Specifically, gaining reading ability should be associated not only with increased word selectivity, but also with decreased selectivity to categories from which word-selective responses are recycled, especially after controlling for age, which often covaries with literacy gains.

Another theory that has been proposed as a driving factor for functional specialization is how that cortical tissue is connected to the rest of the brain, i.e. its connectivity fingerprint (Mahon & Caramazza, 2011). The way that a region is connected will constrain the type of information it has access to from up-stream regions, as well as what information it relays to the rest of the brain (Saygin et al., 2012). This hypothesis has been tested in adults, showing that functional selectivity for different visual categories can be predicted from both structural and functional connectivity (Molloy et al., 2024; Osher et al., 2015). The VWFA, as an input node in a distributed reading network (Vin et al., 2024), has preferential connectivity to the frontotemporal language cortex (Bouhali et al., 2014; Stevens et al., 2017; Yablonski et al., 2024) in adults. Critically, word selectivity that develops after reading acquisition at age 8 can be predicted by the VTC’s structural connectivity with the rest of the brain at age 5 (when no word-selective response was observed) (Saygin et al., 2016). Moreover, recent studies also found that putative VWFA, but not the FFA, preferentially connects to language regions in early childhood (Feng et al., 2022) and even in newborns (J. Li et al., 2020). These prior results support the hypothesis that connectivity precedes and drives functional specialization to some extent. Previous studies also showed that functional connectivity between the VWFA and language regions was associated with reading ability (Y. Li et al., 2017), but this relationship is less clear. For example, Koyama et al. (2011) found a positive relationship between functional connectivity of the left fusiform gyrus and language regions only in adults but not in children, and in a large-scale cross-sectional developmental sample (6-20 years), Yablonski et al. (2024) found that functional connectivity between VWFA and language regions was not correlated with reading ability. Moreover, these studies treat functional connectivity as a static, one-time measurement, making it difficult to dissociate initial connectivity patterns from later developmental changes. Therefore, it is less clear whether VWFA and language connectivity continues to change over development and further contributes to, and/or interacts with, changes and refinement of functional specialization. It is possible that connectivity fingerprints for functional specialization are static and cease to change once they scaffold for a particular functional representation. Alternatively, it is possible that VWFA-Language functional connectivity prior to formal reading education reflects an innate predisposition, whereas subsequent reading experience further strengthens functional connectivity between the VWFA and language regions, contributing to changes in word responses.

Finally, as a high-level visual category-selective region, the VWFA’s tuning profile is primarily visual in nature (J. Li et al., 2024). However, recent studies have also shown that it is sensitive to non-visual linguistic stimuli (J. Li et al., 2024; Planton et al., 2019; Yoncheva et al., 2010). Indeed, long before formal reading instruction, children have extensive exposure to spoken language and the importance of phonological processing in reading acquisition is widely established (e.g., Swanson et al., 2003). However, most studies examining the neural correlates of phonological processing focus on a set of dorsal language or speech regions, such as the left temporo-parietal and inferior frontal cortex, examining activation within or connectivity among these regions (Yu et al., 2018). Very few studies directly examined auditory processing in the VWFA and to our knowledge, no study has examined the development of spoken language sensitivity (if any) within the VWFA. Is the observed high-level linguistic sensitivity in the adult VWFA only a result of years of audiovisual language experience, or does the VWFA already exhibit sensitivity to different aspects of spoken language, either as a precursor to or as a consequence of literacy acquisition?

We attempt to overcome several methodological setbacks that have precluded a clearer understanding of the emergence of word selectivity and the changes that follow literacy. First, by collecting cross-sectional neuroimaging data from prereaders and readers, along with their reading performance, we aim to disentangle the contribution of reading ability from a maturational/general age effect. Second, a precision fMRI approach with multiple functional regions of interest defined within subject helps overcome any spatial blurriness due to group analyses and also helps target the most selective responses by comparing across multiple visual stimulus categories. Third, we used longitudinal data to directly examine the neuronal recycling hypothesis by examining neural tuning prior to and neural changes associated with reading acquisition. Fourth, we additionally collected data during an auditory language task in order to characterize the development of high-level linguistic selectivity within the VTC. Finally, by functionally defining the canonical language network (language-selective temporo-frontal regions), we can investigate the contribution of VWFA-Language connectivity with better precision, rather than relying on coarse estimations of the language cortex using group analysis atlases or anatomical parcellations (J. Li et al., 2020; Saygin et al., 2016; Yu et al., 2022).

In the present study, we collected both cross-sectional and longitudinal data from children in early childhood (3-9 years old). To foreshadow the results, we found that reading acquisition and performance drive the emergence and continued development of word-selectivity in the VWFA; other category-selective cortex is already in place in prereaders, but this category-selectivity increases with age and not with reading performance. Prior to reading acquisition, the pre-VWFA does not show preferential tuning to any particular visual or auditory conditions, and visual word responses increase with continued gains in reading ability without significant decreases in responses to other visual categories. Critically, longitudinal increases in word responses were predicted by changes in functional connectivity to high-level language regions. Finally, readers already showed some sensitivity to auditory language (by a reduction in response to auditory control conditions i.e., texturized language), presumably driven by the observed increase in VWFA connectivity with language regions.

## Methods

### Participants

As part of an ongoing project investigating various cognitive functions in the developing brain, preschoolers and school-age children from the local community of Ohio State University (OSU) were recruited to complete a battery of fMRI and behavioral assessments. A subset of the participants were tested and scanned longitudinally. In total, 79 fMRI sessions (from 44 unique subjects, 37 females) were included in the current study (Figure 1; mean age for prereaders: 4.93 yrs; mean age for readers: 7.10 yrs; age difference: β=2.22, t(68.9)=9.15, p<0.001; see Reading section for prereader/reader information): these participants had completed two runs of the static VWFA localizer and one run of the dynamic visual localizer (see **Methods** below) (at least during their visit at the later time point). Among them, 27 children were scanned multiple times longitudinally; and in the cases where children were scanned more than two times(N=11), sessions from the first and the latest visit with all the data available were used. 22, 27, and 24 participants had the VWFA, language, and dynamic visual localizers at both time points, respectively. Resting-state data were available for 18 of the 27 participants, who were included in the functional connectivity analysis. All children had normal or corrected-to-normal vision and no reported neurological, neuropsychological, or developmental diagnoses at the time of the study (see Table 1 for demographics). The study protocols received approval from the Institutional Review Board at OSU.

**Figure 1.**
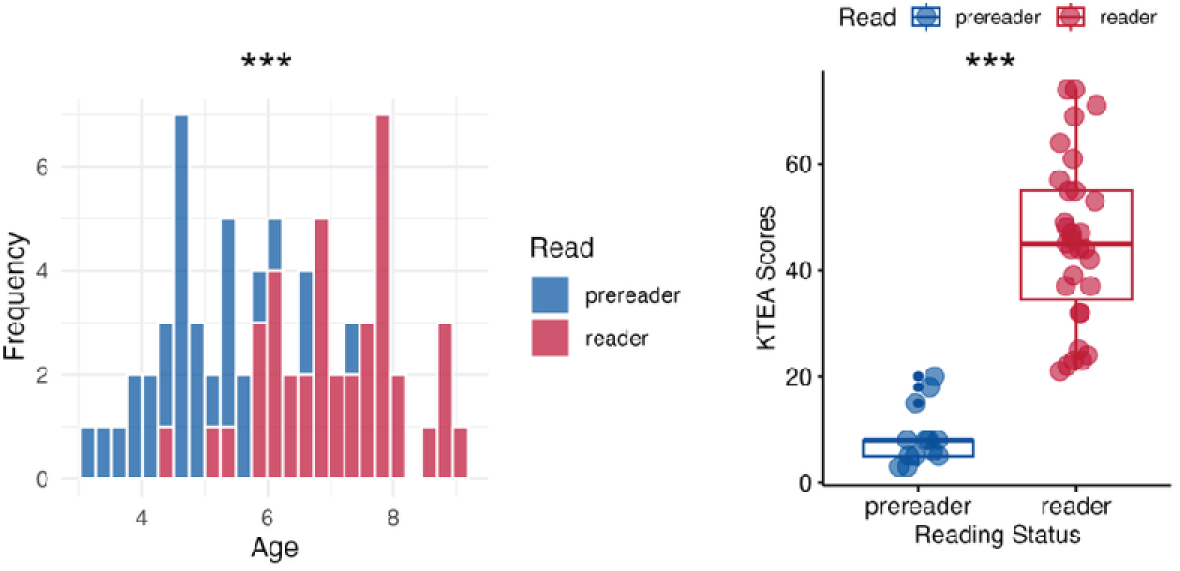
Comparing age and reading ability in prereaders and readers based on the cross-sectional samples. A) Age distribution for prereaders and readers (age difference: β=2.22, t(68.9)=9.15, p<0.001). B) KTEA-3 weighted raw score for prereaders and readers (β=35.68, t(41.6)=7.89, p<0.001).

### Reading

We collected children’s reading status as prereaders or readers based on parents’ reports (“Reading at the time”, yes or no). A subset of the participants also completed the reading comprehension subset of the Kaufman Test of Educational Achievement (Third Edition, KTEA-3). Because not all participants had completed the KTEA-3, we used parent-reported reading status in the main analysis. Note, for participants who had both parent reports and KTEA-3 scores, we relied on the KTEA-3 scores as the criterion: individuals with scores of 20 or below were considered prereaders. Figure 1 below shows the reading comprehension performance from the two groups (reader vs. prereader: β=35.68, t(41.6)=7.89, p<0.001).

### MRI data acquisition and processing

All MRI images were acquired on a Siemens Prisma 3T scanner (at the Center for Cognitive and Behavioral Brain Imaging (CCBBI) at OSU) with a 32-channel phase array receiver head coil.

#### Anatomical

High-resolution anatomical scans were acquired with the whole-head T1-weighted magnetization-prepared rapid acquisition with gradient echo (MPRAGE) sequence (repetition time (TR) = 2300ms, echo time (TE) = 2.9ms, voxel resolution = 1.00 mm^3^). Structural data were then preprocessed by a semi-automated processing stream (FreeSurfer recon-all: https://surfer.nmr.mgh.harvard.edu/fswiki/recon-all) with default parameters. Primary preprocessing steps included motion correction, intensity normalization, skull stripping, white matter and subcortical segmentation, surface reconstruction, and cortical parcellation with the Desikan-Killiany parcellation scheme (Desikan et al., 2006). Critically, a whole brain gray matter mask based on the parcellation was created for each subject and used in the subsequent analyses.

#### Functional

Functional scans were acquired using an echo-planar imaging (EPI) sequence. The majority (60 sessions) of the VWFA localizer data were acquired with the following parameters: TR=2000ms, TE=30ms, base resolution=100×100, voxel size=2 mm^3^ isotropic, 172 TRs. Among them, in 25 sessions, a partial coverage of 25 slices approximately parallel to the base of the temporal lobe to cover the entire ventral temporal cortex was implemented; in 35 additional sessions, a similar protocol with slightly larger voxels (2.2-mm isotropic) was used for whole brain coverage (54 slices). The remaining sessions, as well as the other functional tasks (language and dynamic visual localizers) were acquired with the following protocol: TR=1000ms, TE=28ms, voxel size=2×2×3 mm^3^, base resolution=120×120, 56 slices for whole-brain coverage, and 344 TRs for the VWFA localizer, 244 TRs for the language localizer, 234 TRs for the dynamic localizer. Note that our previous work showed different scanning parameters result in similar results (Li et al., 2024) and all sessions used here had full coverage of the ventral temporal cortex.

Resting-state data were collected across 36 sessions (2 sessions per participant) with the following parameters: TR=1000ms, TE=28ms, voxel size=2×2×3 mm^3^, base resolution=120×120, 56 slices for whole-brain coverage. Sessions were 290 TRs (two sessions were the exception with 580 TRs of data). Most participants were instructed to fixate on the crosshair at the center of the screen (by framing this as a staring contest); 9 participants were shown an animated short film (Partly Cloudy, ∼5 mins) and connectivity was calculated from residuals, see below).

The same preprocessing procedures were implemented for all task data. For motion correction, all time points were aligned to the first time point, and time points with greater than 1 mm total vector motion between consecutive time points were identified to determine whether a participant had too much motion and should be excluded (see below). Other preprocessing steps include distortion correction, detrending, spatial smoothing (3 mm FWHM kernel for the 2 mm^3^ VWFA localizer protocol and resting data, and a 4mm kernel for other tasks). Note that we have previously shown that the effect of smoothing was minimal (Li et al., 2024). Subsequently, the data were aligned from functional to the native anatomical space (see *Registration* below).

For resting-state fMRI, data was first preprocessed with Freesurfer’s FS-FAST preprocessing pipeline (https://surfer.nmr.mgh.harvard.edu/fswiki/preproc-sess) to perform motion correction and also registration to each subject’s native space. Data were further preprocessed using grand mean scaling, smoothing (FWHM = 3 mm), linear interpolation over timepoints with greater than 0.5 mm motion (framewise displacement), and denoising with nuisance regression for the following regressors: mean time course for white matter and cerebral spinal fluid, top 5 temporal principal components of white matter and cerebral spinal fluid separately (calculated using CompCor), and timepoints with greater than 0.5 mm motion (framewise displacement, FD) (Power et al., 2012). For residuals from movie-watching, we took the residual after smoothing with 3mm, linear interpolation over timepoints with greater than 0.5 FD, and modeling time points associated with events in the movie (see Hiersche et al., 2026; Richardson et al., 2018 for more information); these residual images were further denoised as above, using mean time course for white matter and cerebral spinal fluid, top 5 temporal principal components of white matter and cerebral spinal fluid separately (calculated using CompCor). To calculate the functional connectivity between different functional regions, we first averaged the signal from all voxels of a given region and the time courses were correlated with Pearson correlations to estimate the functional connectivity between regions. The resulting correlation values were Fisher’s z-transformed.

### Quality Control

MRI data quality is susceptible to head motion. Several procedures were implemented before, during and after data collection to control and quantify the data quality. First, before scanning, all children were trained using the mock scanner to get familiar with the scanner (e.g., equipment and settings) and listened to recorded audio of scanning noises. Second, foam padding was used to increase comfort for the participants and also fill the space between the head and the coil to reduce the amount of space available for head movement. During the scan, an experimenter stayed in the scanner room to keep the children company; additionally, online motion monitoring allowed the experimenter in the control room to monitor the movement level and signal the experimenter to remind the participant to stay still whenever needed. For anatomical scans, children watched a movie of their choice with an intent to keep them entertained and to minimize motion. After data collection, for each of the functional tasks, functional runs with excessive head motion (more than 25% of time points of a given functional task with > 1 mm total vector motion) were excluded. Participants were excluded if they did not have the minimal amount of data required. Finally, we quantified the amount of motion in each of the functional tasks using the FD values (after removing the outlier volumes). Unsurprisingly, prereaders showed significantly higher motion than readers for the static VWFA localizer task (t(69)=2.86, p=0.006) and the dynamic localizer task (t(69)=2.89, p=0.005), but not for the auditory language task (t(68)=1.85, p=0.067) and resting state (between the two time points: t(17)=0.12, p=0.903). Therefore, whenever we carried out between-subject comparisons, we included FD values of the relevant task(s) as a covariate to control for potential movement effects.

### fMRI tasks

#### VWFA localizer

All participants completed a VWFA localizer task (Saygin et al., 2016). Briefly, printed words (nouns) in black text, scrambled words, line drawings of objects and faces were presented in blocks. All stimuli were displayed on a white rectangular box and a gray grid was superimposed on the stimuli. The scrambled word condition was made by shuffling blocks of this grid of the word condition to preserve low-level visual features of words (lines, curves intersections) (all stimuli are available for download at https://www.zeynepsaygin.com/ZlabResources.html). For each of the visual conditions, 26 stimuli (including 2 repetitions) were presented in a block sequentially for 500ms followed by a 193ms ISI. Each run contains 4 experimental blocks for each of the visual conditions and 3 fixation blocks. The order of these blocks was randomized across participants. A one-back task was implemented to keep participants engaged and two runs of the VWFA localizer task were collected.

#### Dynamic visual localizer

In addition to the static VWFA localizer, we also asked participants to complete a dynamic visual localizer. In this task, video clips (with natural colors) of faces (young children dancing and playing), objects (e.g., moving block toys), bodies (different body parts of kids naturally moving), natural scenes (recorded while driving through suburb areas) and scrambled objects (by scrambling each frame of the object movie clip into a 15-by-15 grid) (Pitcher et al., 2011) were shown and participants were asked to passively watch them. One block consisted of six 3-s video clips from the same category (i.e., 18s per block) and each run consisted of 2 blocks of each experimental condition (alternating in a palindromic manner) with 3 rest blocks with full-screen colors alternating at the beginning, middle and end of each run. The order of blocks was randomized.

#### Language task (auditory)

To assess sensitivity to spoken language, participants also completed an auditory language task (Fedorenko et al., 2010; Hiersche et al., 2024). English sentences (En), nonsense sentences (Ns, which controlled for prosody using phonemically coherent but meaningless words), and texturized speech (Tx, which controlled for low-level auditory features) were presented to participants in blocks through an MRI-safe earphone. Each run included 4 blocks per condition and three 14-second fixation blocks (at the beginning, middle and end). Each block consisted of 3 trials (6 seconds each).

### fMRI analysis

The first-level general linear model was then performed on the preprocessed images: we included experimental conditions for each task as explanatory variables by convolving with the canonical HRF (standard g function, d = 2.25 and t = 1.25) with the block design (standard boxcar function (events on/off)). High motion volumes as well as six motion parameters from the preprocessing stage were included in the first-level GLM as additional nuisance regressors for each task individually. For each run, beta estimates for each condition were extracted for calculating percent signal changes (PSCs, see below). Additionally, we obtained the statistical maps (t statistics) for the contrasts of interest for each task. Specifically, for the VWFA localizer, we obtained contrast maps of words > average of all other conditions, line-drawing objects > average of all others, line-drawing faces > average of all others; similarly, for the dynamic localizer, contrast maps of bodies, scenes, dynamic faces and dynamic objects versus all other conditions were obtained. Finally, for the language localizer, En>Ns, En>Tx and Ns>Tx contrast maps were obtained.

### Define functional regions of interest (fROIs)

We applied the group-constrained subject-specific method (Fedorenko et al., 2010) to individually define fROIs. This method allows us to take the variability of the exact locations of the fROIs into consideration. Previously defined atlases (parcels) that show the typical location of the regions across large samples of adults were used as our search spaces. To investigate the development of the visual category-selective regions, parcels (atlas, created based on the probabilistic maps) from previous studies were used as our search spaces to represent the typical location of the regions across individuals (VWFA, Saygin et al., 2016; FFA, PFS, PPA, and FBA, Julian et al., 2012). In addition, for the longitudinal VWFA-Language connectivity analysis, we also defined frontotemporal language-selective regions. Language functional parcels (2 temporal and 3 frontal) were from Fedorenko et al. (2010) (we used an updated version based on 220 participants, see https://tinyurl.com/5e4tp67w for more details for language parcels). These reference parcels differ in size. For example, compared to the FFA on the right hemisphere, the left FFA (lFFA) was drastically smaller (presumably due to the less reliable face-selective activations on the left and/or more variable locations across individuals). Therefore, we flipped the rFFA to the left, and dilated the smaller parcels (PFS, FFA, FBA, PPA, frontal language parcels) to make the size of parcels comparable.

When defining the fROIs in each individual, we selected the most responsive voxels within a given search space using the statistical maps of the contrasts of interest (see above). The most significant 150 voxels were selected within the reference space and those that showed significant responses to several conditions were assigned to the contrast to which they responded the most. Therefore, it should be noted that even though parcels overlap with each other, especially after dilation due to their close proximity, there is no overlap between the final fROIs -the use of the search space only serves as an initial step to provide a loose guess of the possible location of the fROIs.

### Extracting percent signal change (PSC) and category selectivity

To verify functional selectivity responses and to avoid circularity, we used half of the data (one run of the VWFA localizer and 1/2 run of the dynamic localizer) to define the VTC fROIs and extracted responses within the fROIs from the left-out data. PSC values (calculated based on the raw beta estimates by dividing by the beta values of the baseline condition and multiplying by 100) were reported in the longitudinal analysis to reflect the raw activation of each of the functional conditions. Additionally, we extracted t statistics from the contrasts of interest (based on the first-level GLM modeling) as our selectivity index. This process was repeated, only the data used to define the fROIs and extract responses were switched, and averaged results were reported.

### Registration

A series of registration tools were used to align data across modalities and development (i.e., in longitudinal data). To align functional data to each participant’s native anatomical space, FreeSurfer *bbregister* was used to create the transformation and *mri_vol2vol* was applied to initiate the registration. Note that for exploring category selectivity within the fROIs, all analyses were carried out in this native anatomical space. To align longitudinal data, we computed the affine registration matrix from Freesurfer’s mri_robust_register between the anatomical volumes of the same children scanned at two different time points and initiated the registration with mri_vol2vol. Specifically, we registered the fROIs defined at the later time point to their younger brains to locate the putative VWFA.

### Statistics

To test the presence of category selectivity in the cross-sectional analysis, one-sample t-tests were used to compare whether the selective responses (t statistics from the contrasts of interest) were significantly larger than zero (one-tailed). For longitudinal changes, we also used one-sample t-tests to quantify response changes (against 0, two-tailed) from TP1 to TP2. Multiple comparison correction was carried out using the Holm-Bonferroni method (Holm, 1979) across contrasts or conditions for each region.

Paired t-tests were used to compare selectivity between different contrasts for the cross-sectional analysis. For longitudinal data, one-way repeated ANOVA and follow-up paired t-tests were used to test if the longitudinal fROIs show any functional preference. The results were corrected for the number of pair-wise comparisons.

To test differences between groups, for each contrast, we compared selectivity between readers and prereaders using linear mixed-effects (LME) models with subject as a random intercept and motion parameters (FD values from the two visual localizers) as a covariate. Group differences were tested using estimated marginal means (emmeans) for Read within each contrast, with Holm-corrected p-values across contrasts for each region. In addition, LME models with both KTEA reading scores, age and motion (with subjects as a random intercept) were used to test to what extent the difference between readers and prereaders to their preferred category is uniquely driven by reading ability.

Note that, for analysis in Figure 2, unique participants were used: for children who had multiple time points, we chose the scanning session based on the least motion during the VWFA localizer. When exploring the association between age, reading status & scores, and selectivity, linear mixed-effect models were used to include all available data points (71 sessions) where a random intercept term was added to model the subject effect.

**Figure 2.**
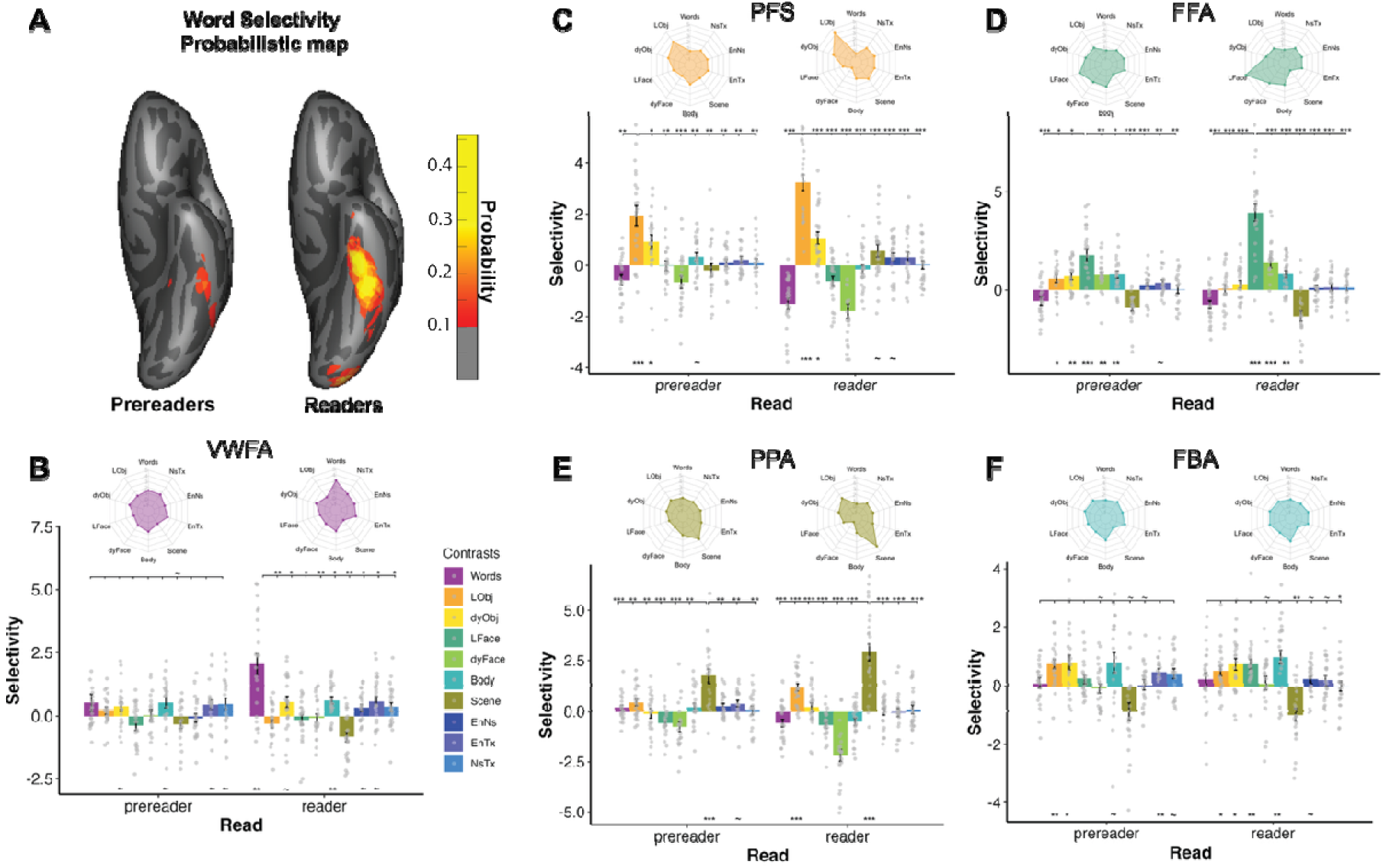
Category selectivity in VWFA and other left VTC fROIs. The selectivity index was the t statistic from the contrasts of interest (from the first-level GLM analysis). For each visual condition, contrasts were created by comparing the conditions of the interests vs. the rest of the conditions in the corresponding task (LObj and LFace, line-drawing objects and faces; dyObj and dyFace, dynamic objects and faces). For auditory language-related contrasts: EnNs, sentences vs. nonsense speech; EnTx, sentences vs. texturized condition; NsTx, nonsense speech vs. texturized. ***, p<0.001, corrected; **, p<0.01, corrected; *, p<0.05, corrected; ∼, p<0.05, uncorrected.

## Results

### Reading acquisition selectively drives the emergence of the word selectivity

We first demonstrated the experience-dependency of the VWFA. Figure 2A shows the probabilistic maps created based on individuals’ word-selective statistic maps (t>2 for individual maps), confirming that at the group level, relatively consistent word selective responses at the left lateral VTC were already observed in readers but not in prereaders. Specifically, as shown in Figure 2B, no word-selective region could be reliably defined in prereaders (word selectivity was not significantly different from 0 (t(21)=1.51, p=0.072). While we failed to identify a reliable VWFA in prereaders, this putative region showed some weak preference for bodies (t(21)=2.46, p=0.011) and objects (dynamic object videos, t(21)=2.11, p=0.023) (however none of these passed multiple comparison correction). Moreover, word selectivity was comparable to objects and bodies (all p>0.1), suggesting no pre-existing selectivity before the development of word selectivity. On the other hand, we confirmed significant word selectivity (t(21)=4.29, p_holm_<0.002) in readers. Additionally, consistent with what we observed in the prereaders, we found weak selectivity for dynamic objects (t(21)=2.23, p=0.018; does not pass correction), and bodies (t(21)=3.51, p_holm_=0.009). Critically, despite these preferences for nonword conditions, the VWFA’s word selectivity (in readers) towered above all other visual category selectivity (word selectivity vs. all other visual selectivity, all p_holm_<0.05) (Table S2). In contrast to the pre-reading putative VWFA, other category-selective regions in the spatial vicinity showed expected selectivity even in the prereaders: we successfully identified PFS, FFA and PPA fROIs in individual subjects that show object, face, and scene selectivity, respectively: their selectivity to the preferred conditions was significantly larger than 0 (Table S1) and also higher than all other non-preferred categories (Figure 2C-E) in both groups (Table S2). FBA, on the other hand, did not show reliable body selectivity in either prereaders or readers (Figure 2F).

Next, we examined selectivity to spoken language within the VTC. Language selectivity was defined by the En>Ns contrast, targeting high-level linguistic information while controlling for speech features such as prosody and rhythm. None of the VTC fROIs showed language selectivity in prereaders. Interestingly, while showing no particular visual selectivity in prereaders, both the putative VWFA and FBA showed some selectivity for meaningless speech sounds (VWFA: Ns>Tx; t(20)=2.00, p=0.029; FBA: t(20)=2.34, p=0.015); however, these did not pass multiple comparison correction. Readers did not show any such trends and instead showed mild language selectivity within the category-selective VWFA (En>Ns, t(21)=2.40, p=0.013) and PFS (En>Ns, t(21)=1.90, p=0.036), as well as in the non-selective FBA (En>Ns, t(21)=1.89, p=0.031). None of these passed multiple comparison correction.

As expected, in the VWFA, readers showed significantly greater word selectivity than prereaders: we modeled the effect of literacy by fitting a linear mixed effect model (including subject as a random intercept and motion as a covariate). Readers showed higher word selectivity than prereaders (β=1.45, t=5.98, p_holm_<0.001). Notably, this significant effect was only observed for word selectivity but not for other category selectivity within the VWFA (all p >0.05; Table S3). Other VTC fROIs also showed greater category selectivity for their preferred categories in readers vs. prereaders (PFS, line-drawing objects: β=0.90, t=3.54, p_holm_=0.004; FFA, line-drawing faces: β=1.24, t=4.97, p_holm_<0.001; PPA, scenes: β=0.97, t=3.95, p_holm_<0.001), except for FBA (β=0.03, t=0.12, p=0.91). Consistent with this, the functional selectivity profiles (Figure 2, neural fingerprint plots) of each region were “pointier” for readers vs. prereaders, suggesting an increase in neural tuning for the preferred category. However, unlike VWFA, the increase in selectivity was also observed for some other non-preferred categories in PFS and PPA (Table S3). Because readers were significantly older than prereaders (see **Method** above), we wanted to disentangle the contribution of literacy acquisition and experience from general maturation effects; we therefore modeled selectivity to the preferred category within each VTC region as a function of both age and reading performance, using the subset of children with available KTEA reading scores. VWFA’s word selectivity was significantly associated with KTEA scores (β=1.14, p=0.030) but not age (β=-0.44, p=0.407) (motion was included as a covariate) (Figure 3A). We also found that the VWFA’s auditory language selectivity (En>Ns) was also positively correlated with reading ability (β=0.44, p=0.041) but not age (β=-0.19, p=0.397) (Figure 3B). Crucially, none of other VTC regions’ selectivity for their preferred categories was significantly associated with reading ability (all p >0.1) (Figure 3C-F). Overall, these results suggest that continued gains in reading ability, rather than maturation, drive the emergence and continued development of word-selectivity and high-level language selectivity within the VWFA.

**Figure 3.**
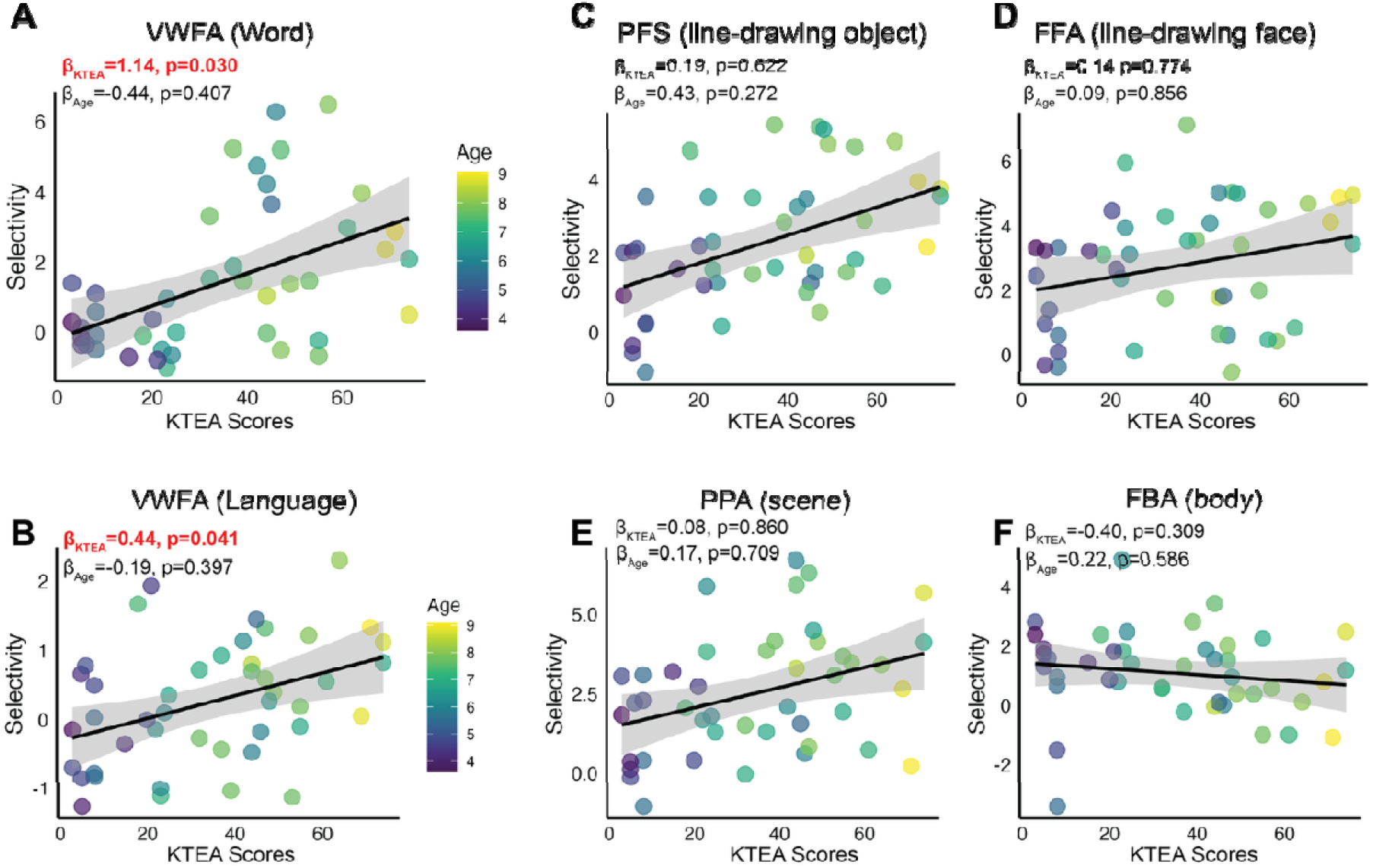
Relationship between category selectivity, age and reading ability. Word, face and object selectivity are from the static visual localizer (A, C-D). Body and scene selectivity are from the dynamic localizer (E-F). Language selectivity (B) is from the auditory language localizer. β estimates are from the linear mixed effect models, including reading scores, age, and motion as the fixed effects, and subject as the random intercept.

On the right hemisphere, no reliable word-selective VWFA could be identified in prereaders (t(21)=0.42, p=0.34); in readers, weak word selectivity was observed, but did not pass correction (t(21)=1.93, p=0.03). Other category-selective regions in the right hemisphere showed expected selective responses to their preferred categories (all p_holm_ <0.01; even for rFBA; see Table S4-5 for details) in both prereaders and readers. Including both KTEA reading score and age in the mixed effect model, we found that only rPFS’s object selectivity to line-drawing of objects was significantly correlated with reading ability (β=0.74, p=0.028) (see Table S6 for details).

### Longitudinal data reveals that the VWFA is not recycled from other category selective cortex

Comparing prereaders vs. readers in cross-sectional data provides evidence for the experience-dependent nature of the VWFA, dissociating it from adjacent selective regions for other visual categories. But to answer the question of neural recycling, we focus on the subset of participants with longitudinal scan sessions and characterize neural signatures in the pre-VWFA prior to developing word selectivity. We identified the VWFA for these children at a later time point (TP2) and projected this region to their earlier time point (TP1) (i.e., pre-VWFA). We then extracted the percent signal changes (PSCs, see Methods) within the pre-VWFA to examine its response profiles to different visual categories, for example, especially those for which neural specialization develops earlier in life (e.g. faces) or those that comprise a majority of a pre-schooler’s visual diet (e.g., limbs and bodies).

One-way repeated-measures ANOVAs showed no main effect of condition (F(3.15,44.16)=1.44, p=0.242) at TP1, suggesting that the pre-VWFA’s responses to various visual and auditory language conditions were generally comparable (Figure 4A). From TP1 to TP2, pre-VWFA’s responses to words increased (responses increased against 0: t(18)=2.07, p=0.054, uncorrected) and critically, at TP2, there was a significant main effect of condition (F(3.42,71.85)=6.66, p<0.001). Post-hoc pairwise t-tests showed that responses to words were significantly higher than responses to all other conditions (all p_holm_<0.05) (Table S7), confirming the cross-sectional results above even with this smaller but more rigorous sample size. Note that as our results showed above, literacy is key to the emergence of the VWFA. Therefore, we divided our longitudinal group into subgroups based on their reading status at each time point: emergent readers (i.e. children who were prereaders at TP1 and readers by TP2) and beginning readers at both time points. As shown in Figure 4B, for emergent readers, a main effect of condition was only found in TP2 (F(10,100)=3.43, p<0.001) but not TP1 (F(10,60)=0.63, p=0.784). Follow-up comparisons indicated that at TP2, the VWFA showed highest responses to words (words vs. dynamic objects, p=0.021; words vs. scenes, p=0.019; words vs. line-objects, line-faces, dynamic faces and bodies, n.s. minimum p=0.097). As for the readers (Figure 4C), a main effect of condition was found for both TP1 (F(10,70)=2.36, p=0.018) and TP2 F(1.9, 19.05)=4.00, p=0.037), with the VWFA showing the highest responses to words at both time points (TP1: all p<0.1, except for words vs. line-faces, p=0.180; TP2, all p<0.1) (Table S7). Altogether these results support the findings in our cross-sectional analyses, and suggest that the emergence of a word-selective VWFA relies on literacy acquisition.

**Figure 4.**
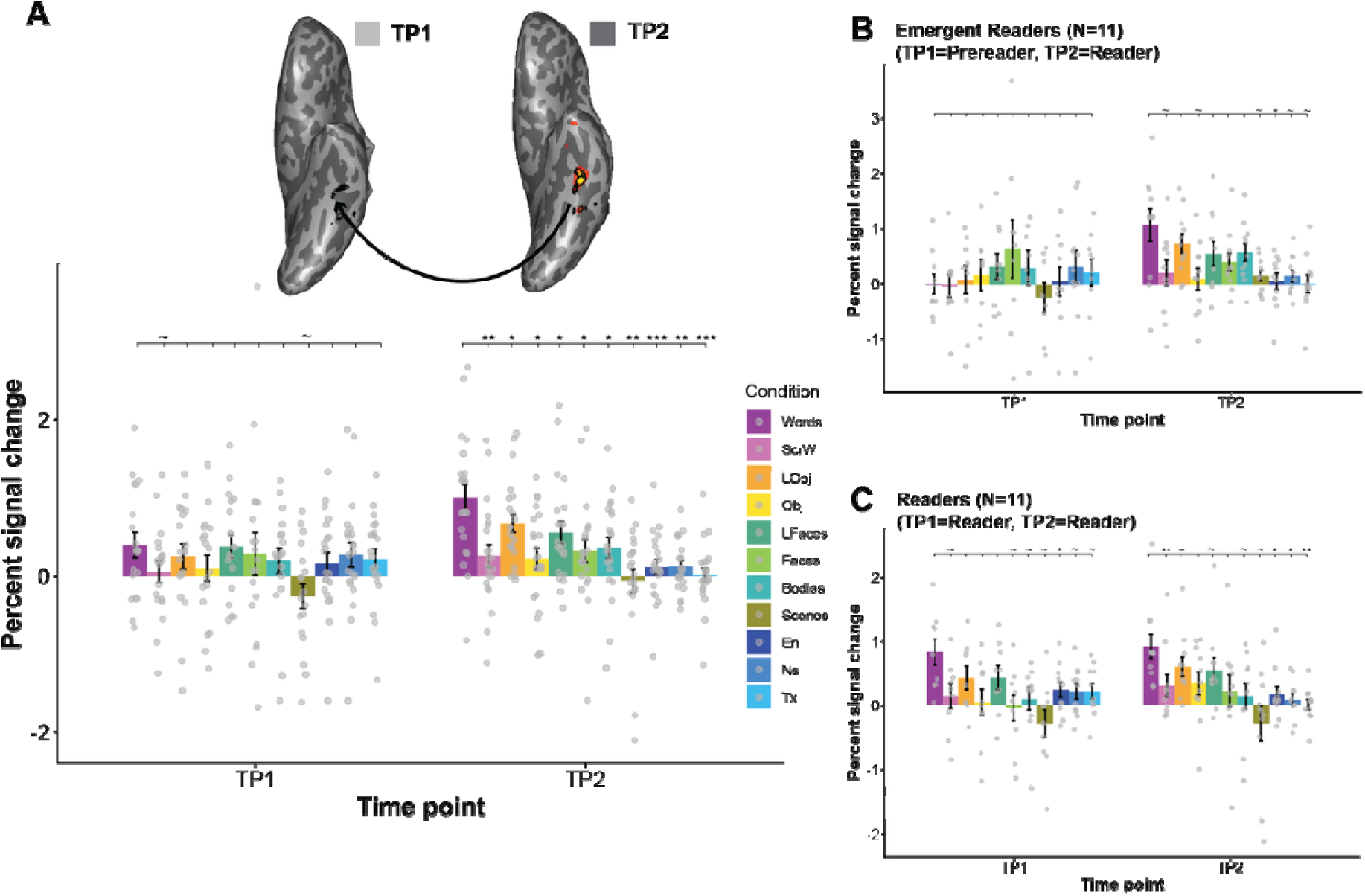
Percent signal change (PSC) in the VWFA for longitudinal samples. A, top, projecting the VWFA defined at the later time point to the earlier brain (pre-VWFA); bottom, PSCs for all visual and auditory language conditions at the earlier (TP1) and later (TP2) time point, respectively. Note that, for TP1, PSCs were extracted from the pre-VWFA. ***, p<0.001, corrected; **, p<0.01, corrected; *, p<0.05, corrected; ∼, p<0.05, uncorrected.

Next, we examined other visual category responses of the VWFA across time to quantify any other changes that may occur as children continue to improve their reading ability. When comparing responses within the VWFA, as shown in Figure 5A, we observed significant increases to line-drawings of objects (t(18)=2.51, p=0.022) (one-sample t-test against 0), but this did not pass correction. None of the changes to the other conditions reached significance (all p >0.19). For other left VTC fROIs, there were no significant changes in activation to any of the preferred and non-preferred high-level visual categories from TP1 to TP2, except that PPA’s responses to bodies significantly decreased (t(18)=-2.66, p=0.016; uncorrected) (Figure 5A). This pattern was also observed when comparing the neural fingerprint plots: as shown in the top panel of Figure 5A, only the VWFA showed visible increases to words and objects, for other regions, their responses remained largely the same from TP1 to TP2.

**Figure 5.**
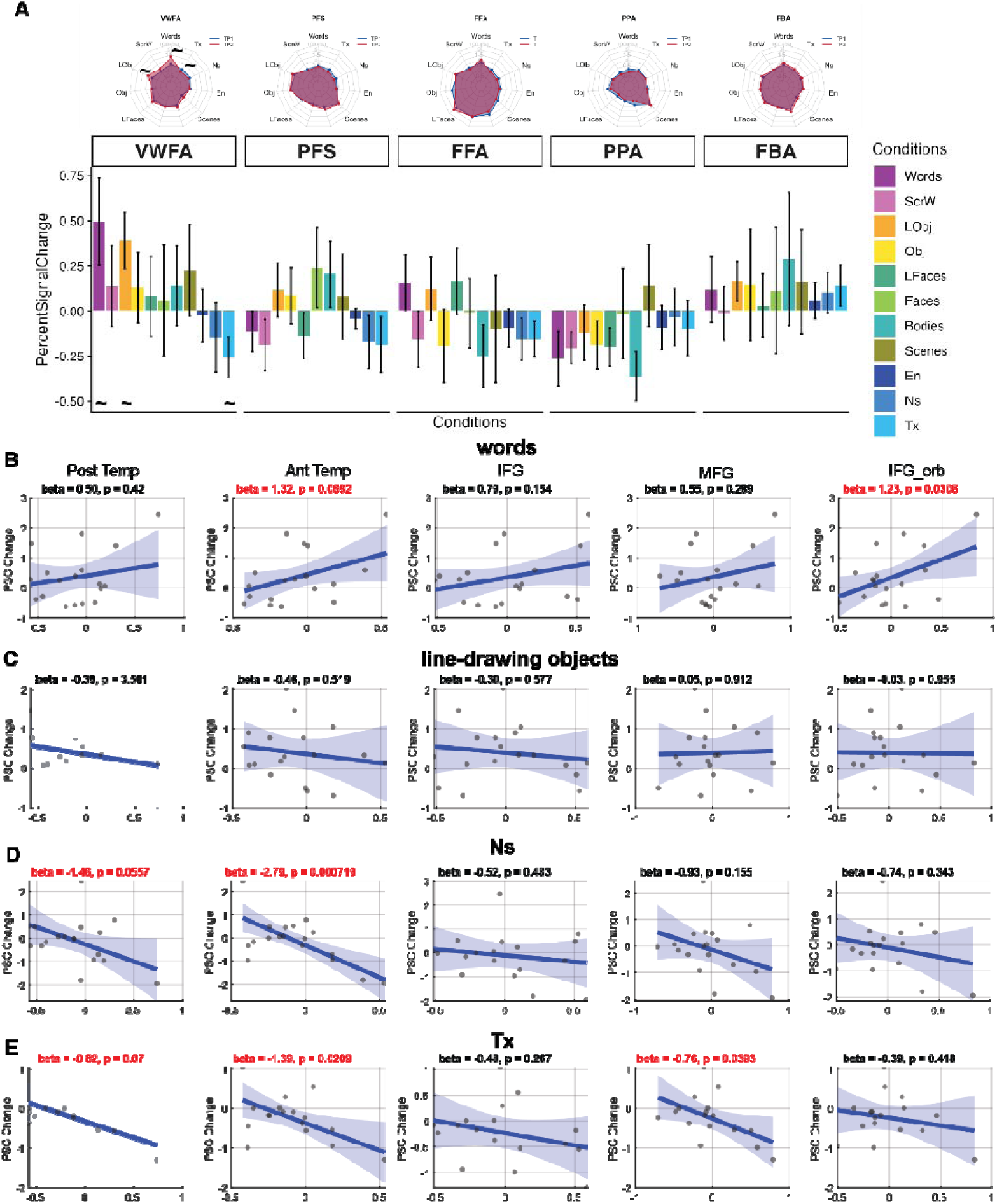
Changes in the VWFA’s functional connectivity to language regions are associated with the changes of the PSCs to words, but not objects, within VWFA. A, top, neural fingerprint plots at TP1 (blue) and TP2 (red) for each fROI; bottom, PSC changes from TP1 to TP2. B-E, correlation between VWFA-Language functional connectivity changes and changes in VWFA’s responses to words (B), line-drawing objects (C), nonsense (D), and texturized sounds (E). Post Temp, posterior temporal; Ant Temp, anterior temporal; IFG, inferior frontal gyrus; IFG_orb, orbital inferior frontal gyrus; MFG, middle frontal gyrus.

We also explored selectivity to auditory language and other auditory conditions in these fROIs across time. While we found no significant selectivity for meaningful auditory sentences as compared to nonsense or texturized sounds at either time point, at TP2, the VWFA, PFS and FFA did start to show higher selectivity for language, mainly by decreases in activation to nonsense and texturized sounds from TP1 to TP2, although only the longitudinal decrease in activation to texturized sounds within the VWFA reached significance (t(20)=-2.31, p=0.032, uncorrected).

Finally, we asked whether the longitudinal response changes within the VWFA were driven by changes in its connectivity with frontotemporal language regions. We found positive correlations between changes in visual word responses of the pre-VWFA’s and changes in its connectivity to anterior temporal (β=1.32, p=0.069) and orbital inferior frontal gyrus (IFG_orb) language regions (β=1.23, p=0.031) (Figure 5B, top). This relationship holds even after controlling for word responses at TP1, suggesting that the increase of functional connectivity with high-level language regions contributes to the increase in word responses regardless of how word-selective it started out (anterior temporal language: β=1.11, p=0.091; IFG_orb: β=0.99, p=0.063). No positive relationship was found between the longitudinal increases in VWFA object responses and its connectivity to language regions (Figure 5B, bottom). Among other category-selective fROIs, only the PFS’s responses to objects were significantly negatively correlated with its connectivity to posterior temporal language fROI, after controlling for TP1 object responses (β=-0.76, p=0.017).

As for the VWFA’s response changes to auditory language conditions, we found that the VWFA’s decreasing response to nonsense and texturized sounds was associated with increased connectivity to language fROIs (Figure 5). After controlling for TP1 responses to these sound conditions, respectively, only connectivity to IFG (β=-0.56, p=0.027) significantly, and connectivity to MFG (β=-0.48, p=0.052) marginally significantly, were associated with decreased responses to texturized sounds.

Altogether, these results suggest that it is unlikely that the VWFA is recycled from regions that are already category-selective to other visual conditions: we only observed an increase in selectivity for words (and line-drawing of objects) in the VWFA but no decreases to other conditions, and other adjacent fROIs maintained their tuning curves for their preferred categories longitudinally. Moreover, we showed that this increase may be driven by the strengthening of its connectivity to frontotemporal language regions.

## Discussion

By combining both cross-sectional and longitudinal data in early childhood, our study investigated factors that drive the emergence and continued development of the VWFA. Confirming the presence of a word-selective VWFA only in readers, but not in prereaders, we also showed that other adjacent category selective cortex was already in place even in prereaders. Moreover, most of this category selectivity continued to increase over time, but only the increase of word selectivity and language selectivity within the VWFA were driven by reading ability but not age. Critically, longitudinal analysis indicated that the pre-VWFA did not show any particular functional tuning for visual or auditory conditions, and the development of word selectivity did not disrupt other non-word category responses within the pre-VWFA. We also observed that the specific neural changes of the VWFA longitudinally across time (increased word responses and decreased responses to nonsense and texturized sounds) were significantly associated with increases in functional connectivity to language-selective regions.

The current study disentangles the role of reading ability and general age-related maturation in the development of category selectivity in the VTC. The emergence of the VWFA in school-age children has been widely studied (Centanni et al., 2017; Dehaene-Lambertz et al., 2018; Nordt et al., 2021; see Chyl et al., 2021 for a review). Meanwhile, reading development occurs concurrently with massive neural and cognitive maturation (e.g., Casey et al., 2005; Gogtay et al., 2004; Toga et al., 2006). To what extent do such developmental changes reflect reading-related increases, rather than broader age-related improvements? Ben-Shachar et al. (2011) found that in addition to age-related increases, changes in word perceptual sensitivity in the left OTC were selectively associated with increases in sighted word efficiency. More recently, Dehaene-Lambertz et al. (2018) found that reading speed correlates with word responses independently of age. However, because word activity in that study was defined relative to rest, it is difficult to conclude that the observed increase was truly word-selective; indeed, a similar relationship was observed between reading speed and face responses. In the current study, by including both age and reading ability in the same model and by using more selective visual contrasts, we provide evidence for the selective contribution of reading ability to VWFA development. Moreover, the response pattern changes between TP1 and TP2 in emergent readers, in comparison with readers (readers at both time points), further support the idea that learning to read gives rise to the VWFA. In addition, utilizing precision fMRI, we also charted the developmental trajectory of other visual selectivity. Consistent with findings that show preferential responses for these categories even in infancy (e.g., Deen et al., 2017; Kamps et al., 2020; Kosakowski et al., 2022, 2024) and differential developmental trajectories for different categories (Scherf et al., 2007; Yan et al., 2024), we found that this early-developed visual selectivity continues to increase. And yet these increases were not associated with reading ability. This result provides further evidence for the functional and spatial specificity of reading-related effects in the emergence of visual word selectivity.

The ubiquitous development of category selectivity within the VTC questions the neuronal recycling hypothesis, which predicts a decrease in existing function alongside the emergence of word selectivity. Our results support a *revised* neuronal recycling hypothesis (Dehaene-Lambertz et al., 2018): pre-VWFA voxels are functionally unspecified and respond similarly to faces, objects, and bodies, and gaining word selectivity did not ‘destroy’ any prior object selectivity (Hervais-Adelman et al., 2019; van Paridon et al., 2021). In fact, the pre-VWFA’s response to line-drawing objects also increased. It is possible that an increase in object responses of the VWFA as a child learns to read could be due to 1) potential semantic representations within the VWFA, given that reading requires linking word forms to meaning, through the semantic/lexical pathway (Coltheart et al., 2001; Plaut et al., 1996; Nation, 2009). This may also explain the positive association between object selectivity and reading ability observed within the rPFS: both reflect gains in semantic knowledge; or 2) the activation of orthographic representations when children internally verbalize nameable objects (orthographic interference) (e.g., Chereau et al., 2007). Interestingly, a previous study showed that the VWFA in struggling readers shows a greater response to objects compared to skilled readers (E. C. Kubota et al., 2019). This finding suggests that, regardless of the processes shared between orthographic and object representations, further functional specialization may be necessary for efficient visual word processing.

Crucially, our study further provided empirical evidence showing that the increase in word responses within the VWFA was associated with changes in its functional connectivity with language-selective fROIs, even after controlling for initial word responses at the earlier time point. This result suggests that the underlying connectivity pattern not only serves as an innate predisposition that initializes the scaffolding of future brain function (Barttfeld et al., 2018; Kamps et al., 2020; J. Li et al., 2020; Saygin et al., 2016), but it also dynamically changes with increasing reading experience. Learning to read requires associating letters with sounds (e.g., Perfetti et al., 1987; Melby-Lervåg et al., 2012) and lexical representations with meanings (e.g., Nation, 2009; Qin et al., 2021). The VWFA may strengthen its connectivity with the frontal and temporal language fROIs with increasing reading experience. Note that most of the existing studies examining VWFA-Language connectivity rely on anatomical approximation for at least one of these nodes. However, these frontotemporal regions house multiple functional networks aside from high-level language that are crucial for reading, such as domain-general cognitive control and semantic cognition (e.g., Binder et al., 2009; Fedorenko et al., 2013). Therefore, by functionally identifying language-selective regions in each individual, our study fills this methodological gap and confirms that it is the VWFA’s connectivity to *language*-selective regions that is associated with the increase in word responses within the VWFA.

In addition to responses to different visual stimuli, the current study was also designed to explore whether there is any sensitivity to spoken language within the VTC in early childhood. In prereaders, we found speech sensitivity, but only within the VTC voxels that were not yet functionally specified (non-selective VWFA and FBA). It is likely that innate connectivity to frontotemporal language regions (Li et al., 2020) already functionally separates these voxels from adjacent areas, allowing these unspecified voxels to show sensitivity to phonological features, and to serve as neural precursors for orthographic selectivity. In line with this idea, previous studies have shown that the onset of VWFA selectivity for visual words coincides with the acquisition of grapheme-to-phoneme correspondences (e.g., Pleisch et al., 2019). Recently, when examining the functional profile of the VWFA in adults, we reported that while it is primarily a visual region, it also shows some sensitivity to high-level linguistic information (Li et al., 2024). The current study further showed that such selectivity emerges in readers (around the age of 7), and that increased functional connectivity with language fROIs may drive the emergence of VWFA language selectivity by decreasing responses to lower-level auditory features in speech. This is consistent with the interactive specialization hypothesis (Johnson, 2001; Johnson et al., 2009) of development, which suggests that functionally-relevant connections increase with increased co-activation (e.g., while reading out loud), leading to increased specialization in connected regions as they continue to interact.

Our study, as well as a few other longitudinal fMRI studies, has provided valuable data on neural tuning of the VTC to visual stimuli (including visual words) (Dehaene-Lambertz et al., 2018; Nordt et al., 2021) over development. However, the resulting picture is far from clear and raises further questions for future research. First, in our study, we found drastic functional profile changes in the prereader-reader group, but less so in the reader-reader group. While the sample size is small after dividing into subgroups and the observed effects were only marginally significant, this suggests that differences in age, reading stage, and the longitudinal sampling window matter. Indeed, inconsistent results in prior literature might be explained by these sample differences: ranging from children with no reading experience who were sampled every two months during the first year of school (Dehaene-Lambertz et al., 2018), to prereading and reading children in early childhood (our study), to children studied later in development (Nordt et al., 2021). Moreover, previous studies have found U-shaped developmental trajectories for word selectivity in older children (Huo et al., 2025; Maurer et al., 2006; Fraga-Gonzalez et al., 2021). Thus, future studies covering not only the period from prereading to the first year of schooling, but also subsequent stages of reading development, ideally with dense sampling intervals (Vinci-Booher et al., 2026), will shed light on potentially nonlinear developmental trajectories for different category selectivity, as well as the different roles that connectivity plays during different stages (E. Kubota, Grill-Spector, et al., 2023; Y. Li et al., 2017; Yablonski et al., 2024).

Second, it is unclear if the size/extent of the VWFA increases with gains in literacy. For our current study, we used a common practice of selecting the top, most selective voxels for a given category (e.g., Fedorenko et al., 2010), and examining its functional selectivity in independent runs. Choosing the most selective voxels of a set size additionally ensures similar sized fROIs, which is important for exploring selectivity and connectivity (see Li et al. 2024 for extensive comparisons showing that different VTC fROI definition methods yield highly similar response profiles in adults). Moreover, we defined longitudinal fROIs and projected back to examine responses at earlier time points (similar to methods in Dehaene-Lambertz et al. (2018)). This approach directly addresses the question of what a cortical tissue does before developing its final functional specialization. However, it is possible that using a hard threshold (e.g., Nordt et al. (2021) and Dehaene-Lambertz et al. (2018)) to define differently-sized fROIs across regions and across age may result in additional information, such as extent of activation (another important measurement of neural development) or overlapping fROIs across regions. These fROIs may be less functionally specified and more susceptible to developmental changes, leaving greater room for plastic adjustments in functional preferences. Indeed, the neural changes reported in Nordt et al. (2021) were observed in developmentally defined emerging/waning regions, rather than in the mature ROI defined at the later time points. Consistent with this idea, studies have found expansion at the periphery of the initial activation of the FFA (Golarai et al., 2007) and that decreases in activity also only found in the periphery (Dehaene et al., 2010). Future work can explore the spatial development of these fROIs in more detail to better understand neural changes associated with gaining literacy.

Third, using both static and dynamic visual localizers, our study echoes the call for moving from only testing 1-2 critical conditions of interest to testing a wider range of visual stimuli to better map out the functional profiles and the development of the VTC regions (e.g., ‘condition rich’ design) (E. Kubota, Grill-Spector, et al., 2023; Ritchie et al., 2026). Indeed, although showing robust selectivity for their preferred categories, each of the VTC regions also responds to other stimuli to some extent (Li et al., 2024). Moreover, compared to adults, the current study seems to yield larger response differences between dynamic and visual localizers for a given category (i.e., faces and objects), further highlighting the importance of the rich-condition approach in shedding light on what visual properties prepare the pre-VWFA to become word selective. For example, it is possible that both limbs and tools share some visual features that serve as a more general organizational principle underlying VTC function, such as spikiness and elongated shapes (Bao et al., 2020) or line junctions (Szwed et al., 2011).

In sum, the current study showed that reading ability uniquely contributes to the emergence and continued development of selectivity for visual words and high-level language within the VWFA and that these increases in selectivity are associated with changes in the VWFA’s functional connectivity with frontal and temporal Language fROIs. This study offers novel insights by disentangling the effects of reading and age, examining mechanisms that contribute to these neural changes, and demonstrating contributing factors that underlie not only visual but also spoken language development within the VWFA.

## Acknowledgement

The authors would like to acknowledge the support from Center for Cognitive and Behavioral Brain Imaging (CCBBI) and Ohio Supercomputer Center (OSC). This material is based upon work supported by the National Science Foundation Graduate Research Fellowship (awarded to K.J.H.) under Grant No. DGE-1343012. Z.M.S. was supported by the Alfred P. Sloan Foundation, OSU’s College of Arts & Sciences, the Chronic Brain Injury initiative at OSU, and OSU’s Women in Philanthropy award and by the National Institutes of Health (NIH) through grant R01HD110401.

## Notes

### Competing Interest Statement

The authors have declared no competing interest.

